# Oil accumulation in leaves driven by a native promoter-gene fusion created using CRISPR/Cas9 mediated genomic deletion

**DOI:** 10.1101/2022.03.08.483411

**Authors:** Rupam Kumar Bhunia, Guillaume Menard, Peter J. Eastmond

## Abstract

Achieving gain-of-function phenotypes without inserting foreign DNA is an important challenge for plant biotechnologists. Here we show that a gene can be brought under the control of a promoter from an upstream gene by deleting the intervening genomic sequence using dual-guide CRISPR/Cas9. We fuse the promoter of a non-essential photosynthesis-related gene to *DIACYLGLYCEROL ACYLTRANSFERASE 2* (*DGAT2*) to drive oil biosynthesis in leaves of a lipase deficient *sugar-dependent 1* mutant. *DGAT2* expression is enhanced more than twenty-fold and the triacylglycerol content increases by around thirty-fold. This strategy offers a transgene-free route to engineering gain-of-function traits such as high lipid forage to increase the productivity and sustainability of ruminant farming.

## Introduction

Transcriptional gain-of-function is a basic tool in plant biotechnology, which traditionally relies on random or targeted insertion of foreign DNA to form a promoter-gene fusion. The introduction of new genetic material has implications since the product is a genetically modified organism (GMO) (1). There are many barriers to commercialisation of GM crops and so it may be desirable to achieve transcriptional gain-of-function by other means, if possible. Genome editing technologies can be used to create single nucleotide substitutions, small insertions, deletions, and rearrangements without foreign DNA integration (2). These changes are widely considered non-GMO (1). It is possible to modulate gene expression by editing *cis*-regulatory elements, but the effects are generally subtle and hard to predict (3). In this study we tested whether the expression pattern of a gene can be radically changed ‘to order’ by bringing it under the control of a promoter from an upstream gene, simply by deleting the intervening genomic sequence using CRISPR/Cas9 (2). Lu et al., (4) recently reported a more complex strategy that relies on large-scale chromosomal inversions and duplications. We engineered leaves to accumulate oil as a proof-of-concept. This is a synthetic trait with the potential to deliver a step change in crop oil yield (5,6), and to increase livestock productivity and suppress enteric methane emissions in pasture-based ruminant farming (7,8).

## Results and discussion

We selected the recipient gene *DIACYLGLYCEROL ACYLTRANSFERASE 2* (*DGAT2*, At3g51520) in *Arabidopsis thaliana*. DGATs synthesise triacylglycerol (TAG) (Fig. 1A) and their overexpression is sufficient to drive ectopic oil production in leaves (9). This oil accumulates particularly when oil turnover is impaired, for example by knocking out the lipase *SUGAR-DEPENDENT1* (*SDP1*) (10) (Fig. 1A). Immediately upstream of *DGAT2* we found a donor gene we named *DGAT2 UPSTREAM GENE 1* (*DUG1*, At3g51510). *DUG1* encodes a chloroplast thylakoid-associated protein of unknown function (11) that is much more strongly expressed in leaves than *DGAT2*, based on public microarray and RNA-Seq data (12). We detected a more than twenty-fold difference in transcript abundance using quantitative RT-PCR (Fig. 1B). We designed gRNA pairs that target within the 5’-untranslated regions (5’-UTRs) of *DUG1* and *DGAT2* (Fig. 1C) and transformed the *sdp1-5* mutant (10) with a dual-guide CRISPR/Cas9 binary vector derived from pEciCAS9-Red (13). We isolated two marker-free homozygous ∼1.6kb genomic deletions, created with different gRNA pairs, following the procedure described by Durr et al., (13), and named them *dug1-1* and *dug1-2* (Fig. 1D & E).

**Figure 1.**
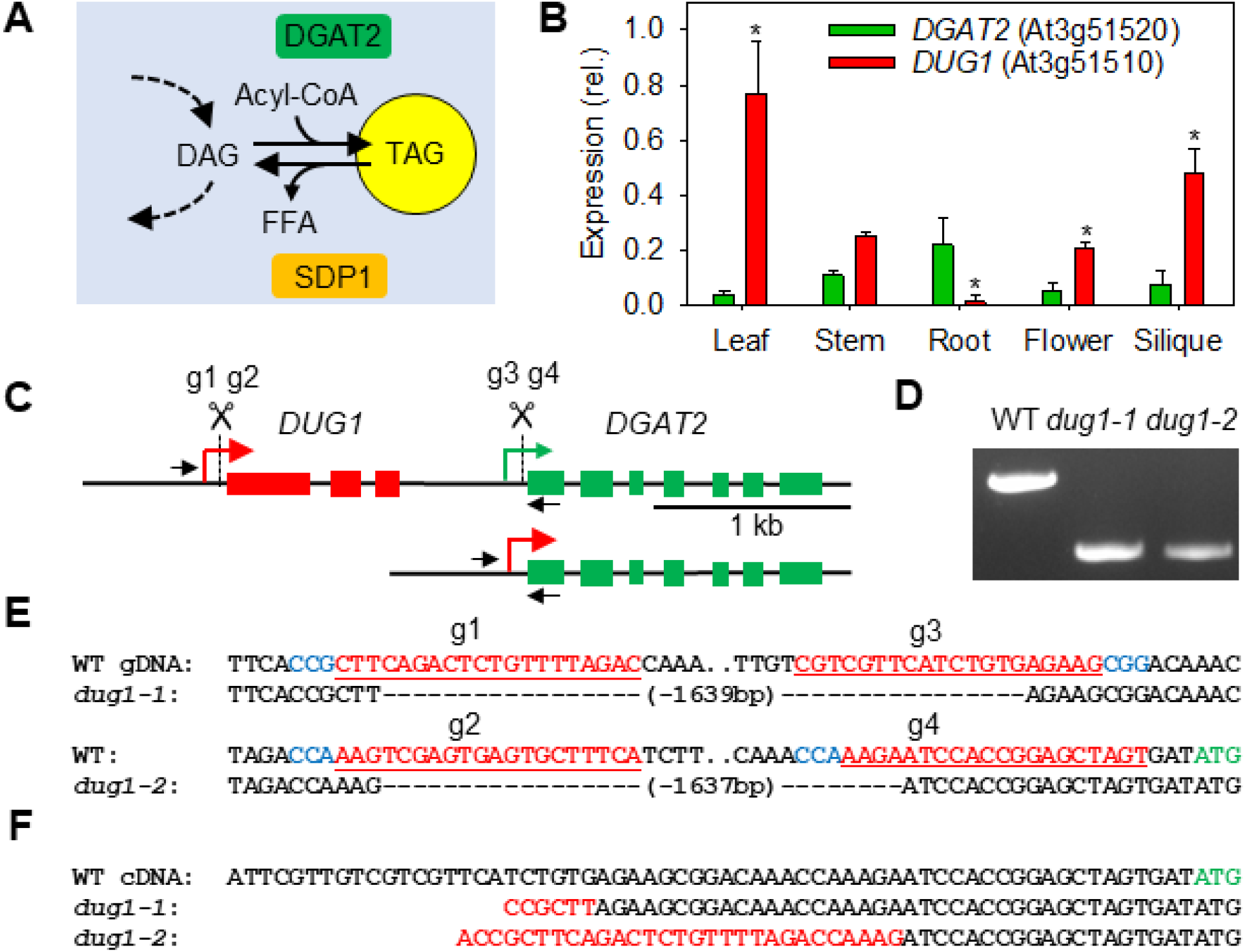
Selection of recipient and donor genes and creation of fusions using CRISPR/Cas9-mediated genomic deletion. (A) Function of recipient gene *DGAT2* (and *SDP1*) in TAG metabolism. DAG, diacylglycerol; Acyl-CoA, fatty acyl-Coenzyme A; FFA, free fatty acid. (B) Quantitative RT-PCR analysis of *DGAT2* and *DUG1* expression in various tissues. Values are presented as mean ± SE (n=3) and are expressed relative to the geometric mean of three reference genes. Asterisks denote values significantly (P < 0.05) different from *DGAT2* (ANOVA + Tukey HSD test). (C) Genomic arrangement of *DGAT2* and *DUG1*. gRNA sites for CRISPR/Cas9 deletion are marked. (D) PCR performed on genomic DNA from homozygous *dug1-1* and *dug1-2* lines. Primer pair are marked on C. (E) Genomic sequence spanning deletion sites. PAMs, gRNA sequences and start codon in blue, red, and green, respectively. (F) 5’-UTRs of *DGAT2* determined by 5’-RACE. *DUG1* sequence in red.

We found that *DGAT2* transcripts possess chimeric 5’-UTRs (Fig. 1F) and their abundance is more than around twenty-fold higher in leaves of *sdp1-5 dug1-1* and *sdp1-5 dug1-2* than in either wild type (WT) or *sdp1-5* (Fig. 2A). Lipid analysis (10) showed that total lipid content of *sdp1-5 dug1-1* and *sdp1-5 dug1-2* leaves is around two-fold higher than in WT or *sdp1-5* (Fig. 2B), and that TAG content is around thirty-fold higher (Fig. 2C). The TAG contains more unsaturated fatty acids (Fig. 2D), consistent with the substrate preference of DGAT2 (9). We also observed the accumulation of lipid droplets within leaf cells by laser scanning confocal microscopy (LSCM) using the fluorescent lipid stain Nile red (10) (Fig. 2E). The rosettes of six weeks old *sdp1-5 dug1-1* and *sdp1-5 dug1-2* plants are marginally smaller than those of either WT or *sdp1-5*, but otherwise appear morphologically normal (Fig. 2F). This suggests that loss of *DUG1* is not critical for chloroplast function.

**Figure 2.**
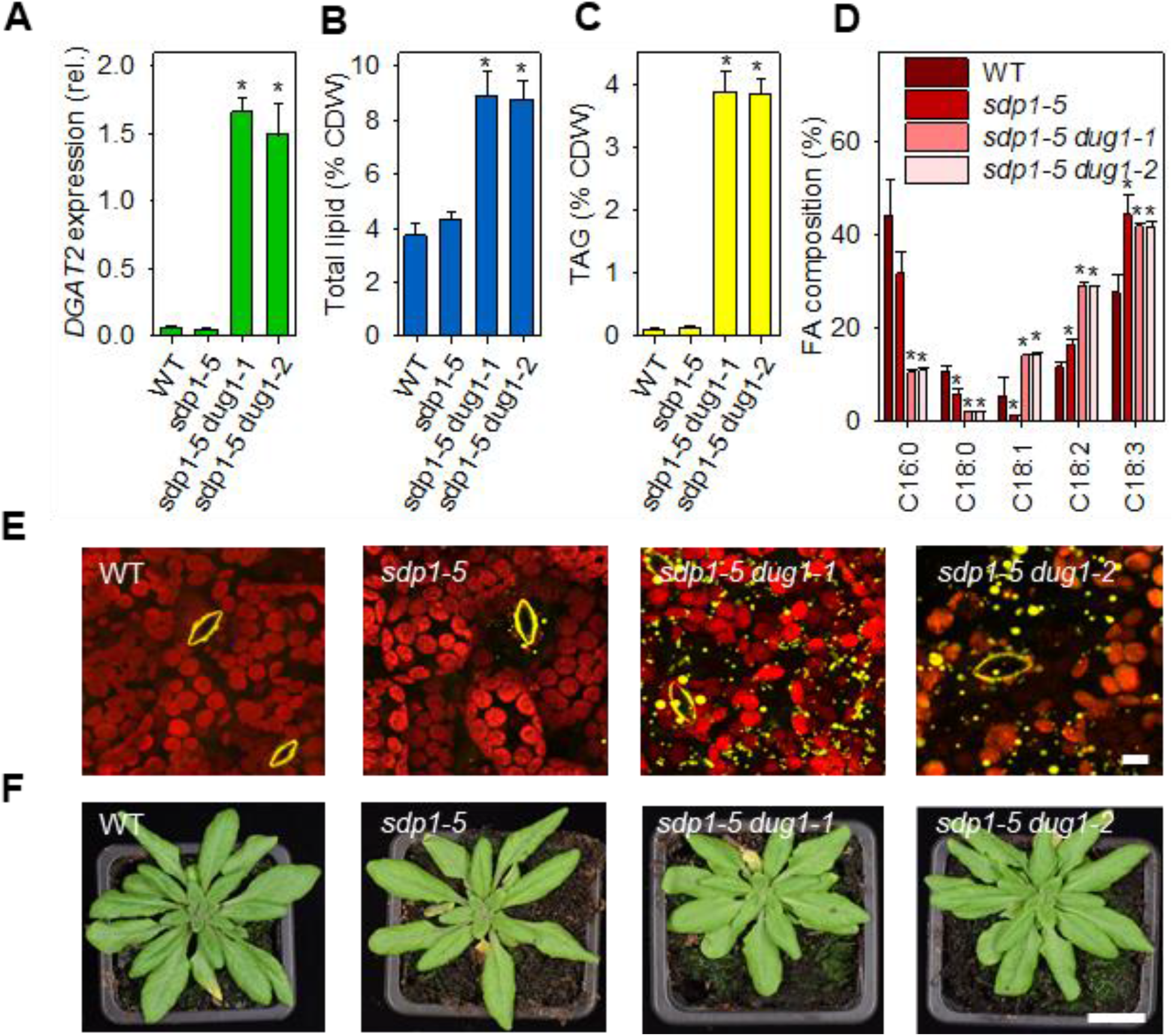
Effect of promoter fusion on lipid metabolism. (A) Expression of *DGAT2* in leaves of *dug1-1* and *dug1-2* in the *sdp1-5* background. (B) Total leaf lipid content and (C) TAG content, as percentage of cell dry weight (CDW). (D) Fatty acid composition of TAG. (E) LSCM images of lipid droplets accumulating in leaves. Lipid droplets and guard cell cuticular ledges yellow (Nile red stained) and chloroplasts red (chlorophyll florescence). (F) Images of rosette plants. In A to D data are presented as mean ± SE (n=3) and asterisks denote values significantly (P < 0.05) different from WT (ANOVA + Tukey HSD test). Scale bars in E & F are 10 µm and 2 cm.

## Conclusions

We show that a tissue-specific transcriptional gain-of-function phenotype can be generated by promoter fusion using CRISPR/Cas9 mediated genomic deletion. This approach relies on an appropriately expressed upstream donor gene lying in the right orientation and within deletion range. Large deletions of tens or even hundreds of kilobases have been achieved in plants using CRISPR/Cas9 (2,13). The approach is also contingent on deletion of the intervening gene(s) being tolerated. In this regard, the strategy could prove most durable in polyploids where gene redundancy is greater (14). Many crop plants and industrial microbial strains are polyploid. Finally, the two-fold increase in leaf total lipid content that we achieve here, without inserting foreign DNA, is likely sufficient to significantly enhance livestock productivity and reduce enteric methane emissions in pasture-based ruminant farming systems, if translated to forage species. Evidence for this has been provided using a GMO approach to enhance oil content in perennial ryegrass (*Lolium perenne*) (7,8).

## Materials and Methods

### Plant material lines and growth conditions

Wild-type *Arabidopsis thaliana* Columbia (Col-0) and *sdp1-5* mutant seeds were described previously (10). For plant growth experiments, the seeds were sterilized, applied to agar plates containing half-strength Murashige and Skoog salts (pH 5.7), and imbibed at 4°C for 4 d. The plates were then transferred to a growth chamber set to 70% relative humidity (16 h light [22°C]/8 h dark [18°C]; PPFD = 250 µmol m^−2^ s^−1^). After two weeks, seedlings were transplanted to 7-cm^2^ pots containing moist Levington F2 compost and the plants were grown on in the growth chamber.

### Cloning and transformation

The CRISPR/Cas9 genome editing method we used to create deletions was adapted from (13). DNA cassettes corresponding to 6728 to 14724 bp of pEciCAS9-Red (GenBank: KY489666) and 52 to 1325 bp of pEN-2xChimera (GenBank: KY489664) were synthesised and cloned into pBinGlyRed and pUC57, respectively. Protospacer sequences were designed to target regions upstream of the transcriptional start site of *DUG1* and *DGAT2* (15) using CRISPR-PLANT (16). They were then synthesised and cloned into pEN-2xChimera using the BpiI and BsmBI restriction enzymes, respectively. The customised gRNAs were then transferred into pEciCAS9-Red by Gateway single-site LR recombination-mediated cloning (13). Arabidopsis plants were transformed via *Agrobacterium tumefaciens*-mediated floral dip (10).

### Selection of germinal deletions

Primary transformant seeds were selected using a Leica M205 fluorescence stereo microscope fitted with a DsRed filter. Plants were grown on soil and genomic DNA was extracted from three-week old plants. For PCR genotyping primers pair DUG1P-F (5’-TGTCGTTTATTTGCACCACG-3’) and DGAT2G-R (5’-AACAGAGAACAAGAGCGACG-3’) were used. Progeny were checked for a 3:1 segregation. DsRed negative seeds of single locus lines were sown on soil and genotyped for deletion events (13). Deletions were confirmed using Sanger sequencing (13).

### Gene expression analysis and 5’-RACE

For each sample, around 100 mg of tissue was ground in liquid nitrogen using a pestle and mortar. The Qiagen Plant RNeasy kit was used to extract RNA and DNase treat it following the manufacturer’s protocol. The Superscript III kit (Invitrogen) was used to produce the cDNA. cDNA samples were normalized, and quantitative PCR was performed on a Roche LightCycler 96 using the FastStart Essential DNA Green Master mix (Roche) with the following condition: preincubation for 10 minutes at 95°C, 45 cycles of three steps amplification (95°C for 10s, 60°C for 15s, 72°C for 15s), melting was performed from 65°C to 97°C. The primer pairs used for *DUG1* and *DGAT2* were QDUG1-F & R (5’-TTCCTCATCCGCTCCG-3’ & 5-CAATGACTCCTGCGGC-3’) and QDGAT2-F & R (5’-TGGTGGAAGCCGGATT-3’ & 5’-CGGGACTTGTGCCTCT-3’). The primers used for the three reference genes (*UBIQUITIN5, ELONGATION FACTOR-1α*, and *ACTIN8*) are described in Bryant et al., (17). Data were analysed using the LighCycler 96 software and Qbase+ (Biogazelle). Analysis of 5’ cDNA ends was performed with the 5’RACE system for Rapid Amplification of cDNA Ends (TermoFisher Scientific) using GSP1 (5’-CCAGGTACAAGAACACAACT-3’) and GSP2 (5’-GAGCAACAACTCCAATCGGTAGCAC-3’).

### Lipid analysis

Total lipids were extracted from homogenized freeze-dried leaf tissue of plants that were six weeks old as described by Kelly et al., (10) and tripentadecanoic acid (15:0 TAG) was added to the homogenized tissue to act as an internal standard. A proportion of the total lipid extract was subjected directly to transmethylation, and the fatty acid methyl esters (FAMEs) were quantified by gas chromatography-flame ionization detection (GC-FID) with reference to the standard (10). The remaining lipid extract was applied to silica thin layer chromatography plates, and neutral lipids were separated using a hexane:diethyether:acetic acid (70:30:1, v/v/v) solvent system. The lipids were visualized under UV light by staining with 0.05% (w/v) primuline in 80% (v/v) acetone, the TAG band was scraped from the plate and transmethylated, and the FAMEs were quantified by GC-FID (10).

### Microscopy

Lipid droplets were imaged in situ by laser scanning confocal microscopy using Nile red staining (10). Nile red stock was made to a concentration of 10 mg mL^−1^ in acetone and diluted to 10 µg mL^−1^ in 0.01% (v/v) Triton x-100 for a working concentration. Detached leaves were vaccume infiltrated with Nile red solution, incubated for 5 h and 1 cm^2^ sections were mounted on slides in water and the abaxial surface imaged with a Zeiss LSM 980 with Airscan 2 (Jena, Germany). The images were captured in SR-8Y mode using the C-Apochromat 40×/1.2 W Korr FCS objective. Nile red was acquired with excitation 0.2% 541 nm Diode-pumped solid state (DPSS) laser and chlorophyll was acquired with 0.4% 639 nm Diode laser. Emission for both channels was 527 – 735 nm to allow fast imaging to match the dynamics of lipid droplet movement. Spectra were obtained from one sample and used for linear unmixing of all samples to eliminate crosstalk between the Nile Red and chlorophyll. Acquisition of Z stacks encompassing the abaxial epidermal and spongey mesophyll cell layers were acquired between 55 and 60µm depth (average Z stack 260 slices), and data are represented as orthogonal XY maximum projections of the Z stack.

## Acknowledgments

We thank Lorraine Berry (Carl Zeiss Ltd.) and Kirstie Halsey for microscopy assistance and Harrie van Erp for transformation assistance. This work was funded by the UK Biotechnology and Biological Sciences Research Council (grant BB/P012663/1). R.K.B. acknowledges support from Rothamsted International and DST-INSPIRE Faculty fellowship schemes.

## Author contributions

P.J.E. conceived the idea and wrote the manuscript. R.K.B., G.M. and P.J.E. conducted the experiments.

